# AMPK-independent LKB1 activity is required for efficient epithelial ovarian cancer metastasis

**DOI:** 10.1101/591073

**Authors:** Adrian Buensuceso, Yudith Ramos Valdes, Gabriel E. DiMattia, Trevor G. Shepherd

## Abstract

Epithelial ovarian cancer (EOC) spreads by direct dissemination of malignant cells and multicellular clusters, known as spheroids, into the peritoneum followed by implantation and growth on abdominal surfaces. Using a spheroid model system of EOC metastasis, we discovered that Liver kinase B1 (LKB1), encoded by the *STK11* gene, and its canonical substrate AMP-activated protein kinase (AMPK) are activated in EOC spheroids, yet only LKB1 is required for cell survival. We have now generated *STK11*-knockout cell lines using normal human FT190 cells and three EOC cell lines, OVCAR8, HeyA8, and iOvCa147. *STK11*KO did not affect growth and viability in adherent culture, but it decreased anchorage-independent growth of EOC cells. EOC spheroids lacking LKB1 had markedly impaired growth and viability, whereas there was no difference in normal FT190 spheroids. To test whether LKB1 loss affects EOC metastasis, we performed intraperitoneal injections of OVCAR8-, HeyA8-, and iOvCa147-*STK11*KO cells, and respective controls. LKB1 loss exhibited a dramatic reduction on tumour burden and metastatic potential; in particular, OVCAR8-*STK11*KO tumours had evidence of extensive necrosis, apoptosis and hypoxia. Interestingly, LKB1 loss did not affect AMPKα phosphorylation in EOC spheroids and tumour xenografts, indicating that LKB1 signaling to support EOC cell survival in spheroids and metastatic tumour growth occurs via other downstream mediators. We identified the dual-specificity phosphatase DUSP4 as a commonly upregulated protein due to LKB1 loss; indeed, *DUSP4* knockdown in HeyA8-*STK11*KO cells restored spheroid formation and viability. Our results strongly indicate that intact LKB1 activity independent of downstream AMPK signaling is required during EOC metastasis.

## INTRODUCTION

Epithelial ovarian cancer (EOC) is a highly lethal malignancy in women since nearly all patients are diagnosed with metastatic disease, at which point the 5-year survival rate is only 30% [1]. Standard-of-care for late-stage EOC has largely unchanged over 25 years and encompasses aggressive cytoreductive surgery with combination carboplatin/paclitaxel chemotherapy. Most patients initially respond to treatment, but recurrence and chemo-resistance is common and fatal in most cases. Therefore, the development of novel approaches to impede metastasis and recurrence of chemo-resistant disease will be paramount to improve outcomes for women diagnosed with EOC [2].

A common symptom of advanced EOC is the accumulation of ascites within the peritoneal cavity [3]. This ascites often harbors individual malignant cells or those existing as multicellular aggregates, herein referred to as spheroids, the latter of which offer numerous selective advantages in the context of metastatic dissemination [4]. EOC spheroids exhibit anoikis resistance and possess an enhanced capacity to adhere to ECM and displace mesothelial cell monolayers [3; 5; 6]. In addition, recent evidence indicates that EOC spheroids seed peritoneal metastases that maintain the cellular heterogeneity of the primary tumour [7]. Thus, EOC spheroids likely play a key role in efficient intraperitoneal metastasis.

We previously reported that Liver kinase B1 (LKB1; encoded by the *STK11* gene) is expressed in the majority of EOC cell lines and patient ascites-derived cells; importantly, transient knockdown of *STK11* in EOC cells results in reduced spheroid viability and increased chemo-sensitization [8]. LKB1 is a ubiquitously-expressed, highly-conserved serine-threonine kinase that commonly acts as the upstream kinase for AMP-activated protein kinase (AMPK), primarily in regulating cell metabolism [9]. The LKB1-AMPK signaling axis functions as a sensor of energy status in the cell and enables adaptation to conditions that deplete intracellular ATP. Decreased ATP and the corresponding increase in AMP favors phosphorylation and activation of AMPK by LKB1, resulting in the coordinated downregulation of anabolic pathways and upregulation of catabolic pathways to restore energy homeostasis [10].

LKB1 is often described as having tumour suppressor-like activity since specific cancers feature somatic alterations in the LKB1-encoding gene *STK11* [11–13] or have decreased LKB1 protein expression [14; 15]. In addition, germline inactivating mutations in *STK11* have been implicated in Peutz-Jeghers syndrome (PJS) [16], which is characterized by the formation of benign polyps in the gastrointestinal tract and an increased risk for several cancers [17]. LKB1 loss has also been implicated in EOC initiation [15; 18]; however, there is significant evidence suggesting that LKB1 and its direct signaling partners function in context-dependent, pro-tumorigenic capacities [8; 19-22].

To date, no studies have assessed the impact of LKB1 loss-of-function in mouse models of cancer metastasis, such as late-stage EOC. In this study, we sought to determine how complete LKB1 ablation in EOC cells would affect metastatic potential. We present evidence indicating that LKB1 loss decreases anchorage-independent viability of EOC cells and reduces the long-term spheroid maintenance. In a similar fashion, LKB1 loss in three independent EOC cell lines significantly reduced tumour burden and extended host survival in a xenograft mouse model of peritoneal metastasis. Based on these findings, we propose that strategies targeting LKB1 or its signaling partners would have therapeutic potential in metastatic EOC.

## RESULTS

### LKB1 lacks tumour suppressor-like activity in EOC

We previously reported that LKB1 protein is expressed in EOC tumour samples, and that transient *STK11* knockdown decreases EOC spheroid viability and chemo-resistance, suggesting that LKB1 or its signaling partners might represent viable therapeutic targets in EOC [8]. Therefore, we sought to investigate how sustained LKB1 loss would affect tumourigenic and metastatic potential in several different EOC cell lines. OVCAR8 (high-grade serous ovarian cancer; HGSOC) and HeyA8 (poorly-differentiated ovarian cancer) cells form robust spheroids as well as aggressive tumour growth in immune-compromised mice, and therefore represent two EOC cell lines with which to test our hypothesis regarding LKB1 function. The iOvCa147 cell line was derived by our group from the ascites of a heavily-treated HGSOC patient [23]. This cell line forms less dense yet viable spheroids, and establishes ascites and tumours in NOD/SCID mice with longer latency as compared with OVCAR8 and HeyA8 cells. We used CRISPR/Cas9-based genome editing to generate stable cell lines lacking LKB1 expression: OVCAR8-*STK11*KO, HeyA8-*STK11*KO, and iOvCa147-*STK11*KO, as well as one immortalized human fallopian tube secretory epithelial cell line (FT190-*STK11*KO) for comparison. Complete ablation of LKB1 expression was confirmed in each *STK11*KO cell line (**Figure 1A**). Under standard adherent culture conditions, *STK11*KO cell lines resembled parental controls with respect to morphology; iOvCa147-*STK11*KO cells were the notable exception and had a spindle-shaped appearance compared to controls **(Supplemental Figure S1)**. Stable LKB1 loss in our newly-derived *STK11*KO cell lines did not affect proliferation since doubling time was unchanged compared with parental cell lines (**Figure 1B**).

**Figure 1.**
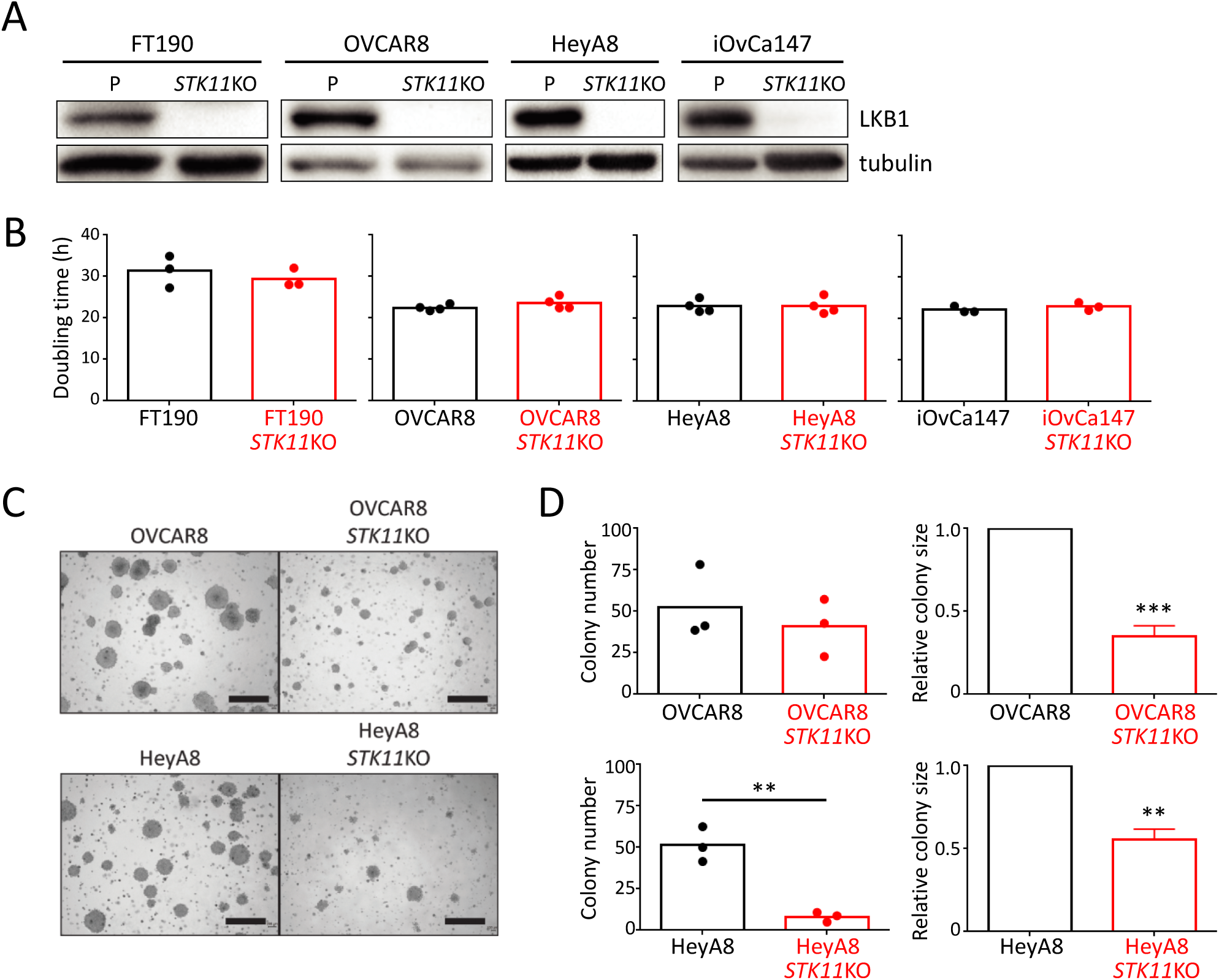
Loss of LKB1 reduces EOC cell anchorage-independent growth. **(A)** Immunoblot analysis of FTE and EOC *STK11*KO cell lines and matched parental controls to confirm loss of LKB1 protein expression in CRISPR/Cas9-edited lines. **(B)** Doubling time for parental and *STK11*KO cell lines in adherent culture conditions. Confluence was monitored over 3 days in adherent culture using the IncuCyte Zoom imaging system. Doubling time was determined using GraphPad Prism 6.0 by fitting an exponential growth curve to confluence over time data in at least 3 independent experiments. **(C)** Images of OVCAR8, OVCAR8-*STK11*KO, HeyA8 and HeyA8-*STK11*KO cell lines cultured in soft agar to assess anchorage-independent colony forming potential. Images are representative of 3 independent experiments. Scale bars represent 500 µm. **(D)** Quantification of soft agar colony formation. Data are presented as colony counts per field of view and relative colony size within each independent experiment. Groups were compared using the two-tailed Student’s *t*-test (**, *p* < 0.01; ***, *p* < 0.001).

It is well-established that the majority of HGSOC likely originate in the secretory epithelium of the distal fimbriae of the fallopian tube [24–26], and mouse models also support this as a primary cell-of-origin in this disease [27–29]. Two reports have implied that loss of LKB1 expression may be involved in the development of HGSOC [15; 18] whereas we have alternative conflicting data demonstrating that LKB1 is in fact expressed and functional in late-stage EOC [8]. If LKB1 has tumour-suppressive properties in EOC, then *STK11* knockout should enhance normal FT190 cell clonogenicity in both anchorage-dependent and -independent conditions. We performed clonogenic assays using each line harboring LKB1 loss to determine whether anchorage-dependent replicative potential would be affected by loss of LKB1 in FTE and EOC cells. We found that LKB1 loss significantly decreased clonogenic potential in FT190 cells, but not in any EOC cell lines tested **(Supplemental Figure S2**).

Since anchorage-independence is a common feature of transformed cells and critical to EOC metastasis, we determined how LKB1 loss would affect anchorage-independence in normal FTE and transformed EOC cells. In contrast to the results from adherent clonogenic assays, loss of LKB1 significantly decreased soft agar colony number and colony size in HeyA8 cells (**Figure 1C & D**); OVCAR8-*STK11*KO soft agar colony number was not different from parental controls, however colony size was significantly decreased (**Figure 1C & D**). The iOvCa147 cell line has a very poor colony-forming capacity in soft agar and were thus not tested by this assay. Furthermore, FT190 cells, which have been immortalized due to p53 and pRB inactivation by SV40 large T antigen [30], were unable to establish any colonies in soft agar with or without intact LKB1 (data not shown). These results suggest that LKB1 does not possess overt tumour suppressive activity, but may be required for aggressive EOC cell growth or viability when in suspension.

### LKB1 is required in EOC spheroids to maintain cell viability

Spheroids are frequently present in the peritoneal ascites of EOC patients, and facilitate peritoneal metastasis since they exhibit anoikis resistance, adhere to mesothelial cell monolayers [3; 5], and help to establish peritoneal metastases [7]. We have demonstrated previously that decreased LKB1 expression by transient *STK11* knockdown has little to no effect on the growth and viability of adherent proliferating EOC cells, yet viability is reduced in spheroids [8]. Thus, we sought to determine how sustained loss of LKB1 would affect FTE and EOC spheroid viability. We cultured FTE and EOC cells in ultra-low attachment (ULA) culture plates to force cells into suspension (**Figure 2A**). LKB1 loss significantly decreased the number of viable cells for all EOC cell lines tested (**Figure 2B**), and the proportion of dead iOvCa147-*STK11*KO cells significantly increased, indicating that LKB1 is required for maximal EOC spheroid viability in this *in vitro* model of metastasis. Normal FT190 cells exhibited rapid attrition in spheroid culture (**Figure 2A**) with even further loss in cell viability due to LKB1 loss (**Figure 2B**), indicating again that LKB1 loss does not possess tumour suppressor-like activity when FTE cells are deprived of attachment to a substratum. Long-term spheroid culture of HeyA8 and OVCAR8 cells yielded an overt decrease in growth and viability due to LKB1 loss (**Supplementary Videos**), and this was especially pronounced when spheroids were maintained for up to 21 days (**Figure 2C**). This latter result provided strong *in vitro* evidence suggesting that tumour-forming and metastatic potential of EOC cells *in vivo* may be blunted by loss of LKB1.

**Figure 2.**
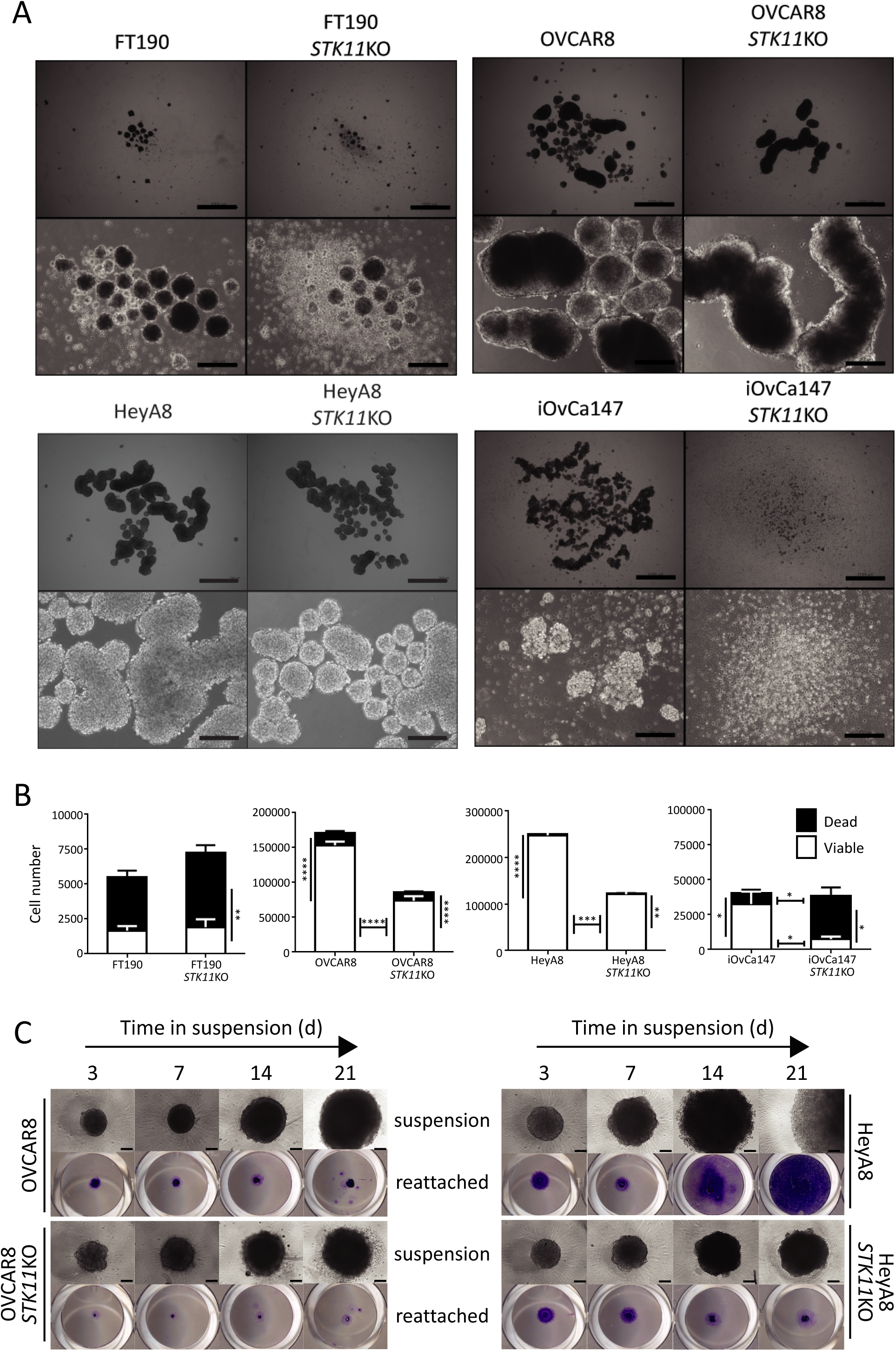
LKB1 is required for EOC spheroid cell viability. **(A)** Images of FTE and EOC cell lines cultured in suspension using ultra-low attachment (ULA) vessels. Images were captured at the following time points (post-seeding): FT190, FT190-*STK11*KO, day 7; OVCAR8, OVCAR8-*STK11*KO, day 5; HeyA8, HeyA8-*STK11*KO, iOvCa147, iOvCa147-*STK11*KO, day 3. Scale bars represent 1 mm (upper panels) and 250 µm (lower panels). **(B)** Cell viability for FTE and EOC cell lines cultured in suspension using ULA vessels. Viable and non-viable cell numbers were determined by trypan blue exclusion cell counting in a minimum of 3 independent experiments. Data are presented as the number of dead and viable cells per well of a 24-well plate (mean ± SEM). Two-way ANOVA and Tukey’s multiple comparisons test were performed for statistical analysis (*, *p* < 0.05; **, *p* < 0.01; ***, *p* < 0.001; ****, *p* < 0.0001). (C) LKB1 is required for long-term growth and viability of EOC spheroids. EOC cells were cultured in 96-well round-bottom ULA plates to form single spheroids for the indicated time periods up to 21 days and phase contrast images were captured (upper panels, “suspension”). At each indicated time point, individual spheroids were transferred to single wells of a 48-well adherent culture plate for 1 day (HeyA8, HeyA8-*STK11*KO) or 2 days (OVCAR8, OVCAR8-*STK11*KO) for sufficient re-attachment. Following transfer of spheroids into adherent culture, cells were fixed and stained (lower panels, “reattached”). Scale bars represent 200 µm.

### LKB1 is required to maintain EOC metastatic potential

The results of our *in vitro* studies indicate that LKB1 is required for both anchorage-independence and spheroid viability in EOC cells. Since spheroids play an important role in EOC metastasis, we sought to determine the effect of LKB1 loss in an *in vivo* xenograft model of peritoneal metastasis. EOC cells are injected directly into the peritoneal space of immunodeficient host mice to mimic metastasis, where tumour pathology and pattern of dissemination are similar to late-stage disease in patients [31]. We hypothesized that stable LKB1 ablation in EOC cells would decrease metastatic potential *in vivo*.

We found that survival was extended in hosts for all three *STK11*KO cell lines when compared with parental cell lines (**Figure 3A**). Compared to parental control hosts, median survival was 48.5% longer for OVCAR8-*STK11*KO hosts (78 d *versus* 52 d; log-rank test, *p* = 0.0005), but only 17.1% longer for HeyA8-*STK11*KO hosts, which approached statistical significance (51 d *versus* 41 d; log-rank test, *p* = 0.1105). Median survival for iOvCa147-*STK11*KO hosts was undetermined since most mice survived until study completion, yet survival was significantly longer than the parental control group (log-rank test, *p* = 0.0015).

**Figure 3.**
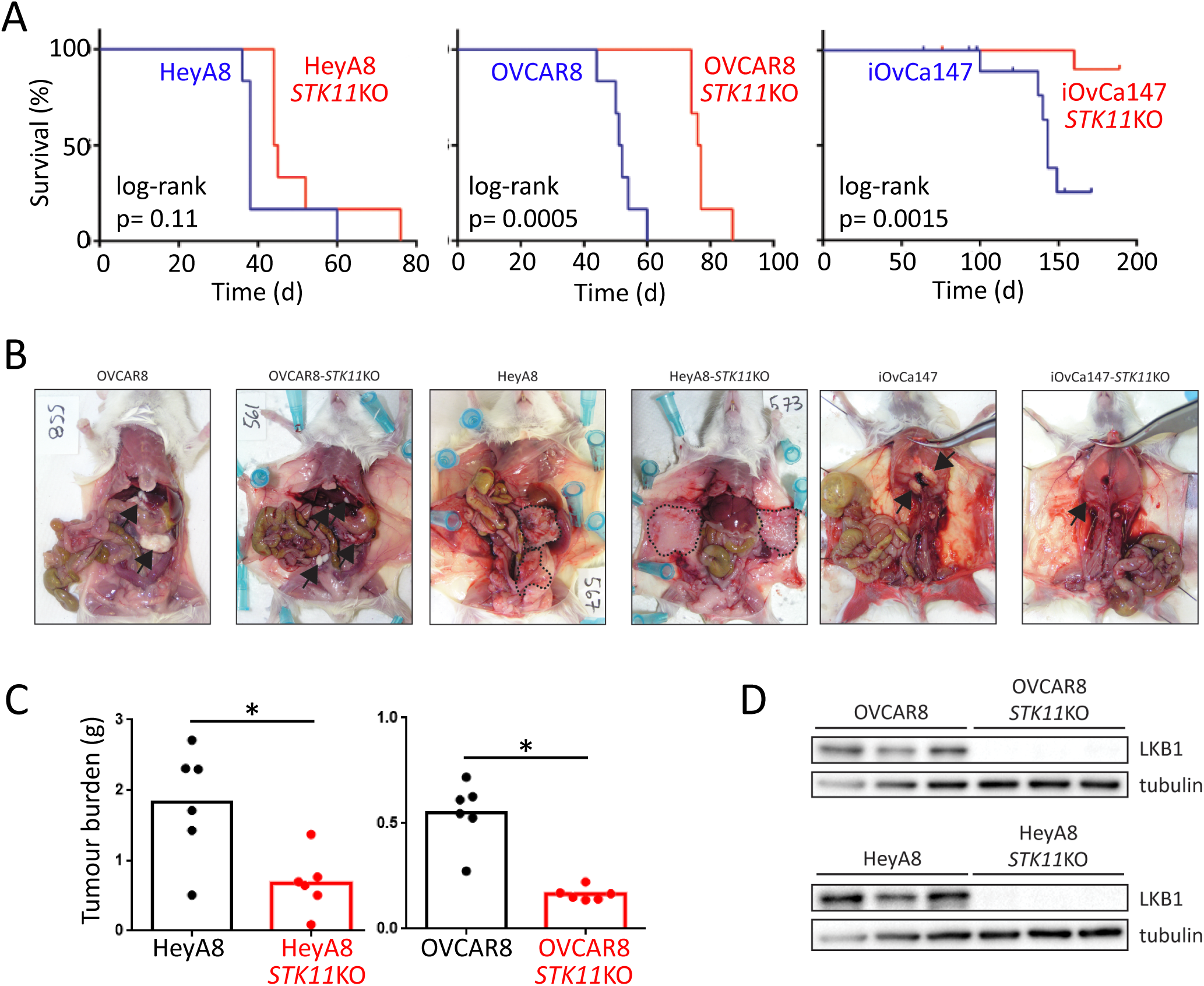
EOC cells lacking LKB1 have decreased metastatic potential. **(A)** Survival analysis for parental and *STK11*KO EOC cell lines. EOC cells were injected i.p. into female NOD/SCID mice. Mice were euthanized when humane endpoint criteria were reached. Censored subjects due to thymic lymphoma in the iOvCa147/iOvCa147-*STK11*KO study are indicated by tick marks on the survival curve. OVCAR8, OVCAR8-*STK11*KO, HeyA8, HeyA8-*STK11*KO, n=6 per group; iOvCa147, iOvCa147-*STK11*KO, n=12 per group. Parental and *STK11*KO survival curves were compared using the log-rank test. **(B)** Representative images of mice at time of necropsy. Tumours are indicated by black arrows (OVCAR8 and iOvCa147) or enclosed within a dotted black boundary (HeyA8). **(C)** Quantification of total tumour burden. Mice were euthanized at a single time point post-injection (OVCAR8, OVCAR8-*STK11*KO, 51d; HeyA8, HeyA8-*STK11*KO, 33d) and all macroscopically visible tumours were excised and weighed together. Data are presented as the total tumour wet weight per mouse and the overall mean. Parental and *STK11*KO groups were compared using the two-tailed Student’s *t*-test (*, *p* < 0.05). **(D)** Immunoblot analysis on excised tumours to confirm absence of LKB1 in OVCAR8- and HeyA8-*STK11*KO tumours. Tubulin was used as a loading control. Each lane represents a tumour from a different mouse.

Frequency of ascites accumulation was decreased in OVCAR8-*STK11*KO relative to OVCAR8 injected mice, but overall dissemination patterns were similar, indicating that metastatic trajectory was unchanged by LKB1 loss (**Figure 3B and Supplementary Table S1)**. Macroscopic tumour burden in OVCAR8-*STK11*KO injected hosts was significantly decreased compared to OVCAR8 cells (**Figure 3B**). In mice injected with HeyA8-*STK11*KO cells and HeyA8 controls, we observed an altered pattern of tumour dissemination associated with LKB1 loss. HeyA8-*STK11*KO-injected hosts exhibited a thin layer of tumour cells adherent to the peritoneal wall, and little evidence of solid tumour lesions at other sites (**Figure 3B and Supplementary Table S1)**. In contrast, HeyA8-injected hosts had several large and solitary solid tumour masses at the omentum and lower peritoneal cavity, often adherent to the intestine and uterus (**Figure 3B and Supplementary Table S1)**. This resulted in a total tumour burden that was significantly reduced in HeyA8-*STK11*KO injected mice as compared with HeyA8 controls (**Figure 3C**). Sustained LKB1 loss was confirmed by immunoblotting of protein lysates from excised OVCAR8- and HeyA8-*STK11*KO tumours (**Figure 3D**).

Similar to our *in vitro* spheroid assays, we observed the most striking effects of LKB1 loss in the iOvCa147 cell line, which was originally generated from the ascites of a high-grade serous ovarian cancer patient by our group [23]. In line with the extended survival of iOvCa147-*STK11*KO hosts (**Figure 3A**), peritoneal colonization was substantially diminished, with most mice exhibiting little evidence of macroscopically-observable peritoneal tumours upon necropsy (**Figure 3B**). In contrast to iOvCa147-*STK11*KO hosts, mice injected with iOvCa147 parental control cells exhibited peritoneal tumours at multiple sites, with particularly frequent colonization at the omentum, peritoneal wall and diaphragm. Since most iOvCa147-*STK11*KO mice failed to develop tumours, we did not measure overall tumour burden in this cohort.

Histological analysis indicated that OVCAR8-*STK11*KO tumour nodules frequently contained large necrotic regions, and immunohistological staining revealed that Ki-67-positive cells were frequently restricted to the periphery of tumour nodules with a core of cleaved caspase 3-positive apoptotic cells (**Figure 4A**). Compared to parental OVCAR8 controls, these highly-necrotic/apoptotic OVCAR8-*STK11*KO tumours had increased carbonic anhydrase 9 protein expression **(Figures 4B & C)**, which is indicative of elevated hypoxia in tumours [32]. In contrast, enhanced necrosis and apoptosis was not observed in HeyA8-*STK11*KO tumour xenografts, however there appeared to be fewer Ki-67 positive cells compared with HeyA8 tumours (**Figure 4A**). Very few iOvCa147-*STK11*KO hosts developed any tumours (**Figure 3A and Supplementary Table S1**). However, histologic analysis of a single iOvCa147-*STK11*KO lesion revealed thinly-deposited EOC cells with papillary features that were not evident in the solid tumours arising in iOvCa147 controls (**Figure 4A**).

**Figure 4.**
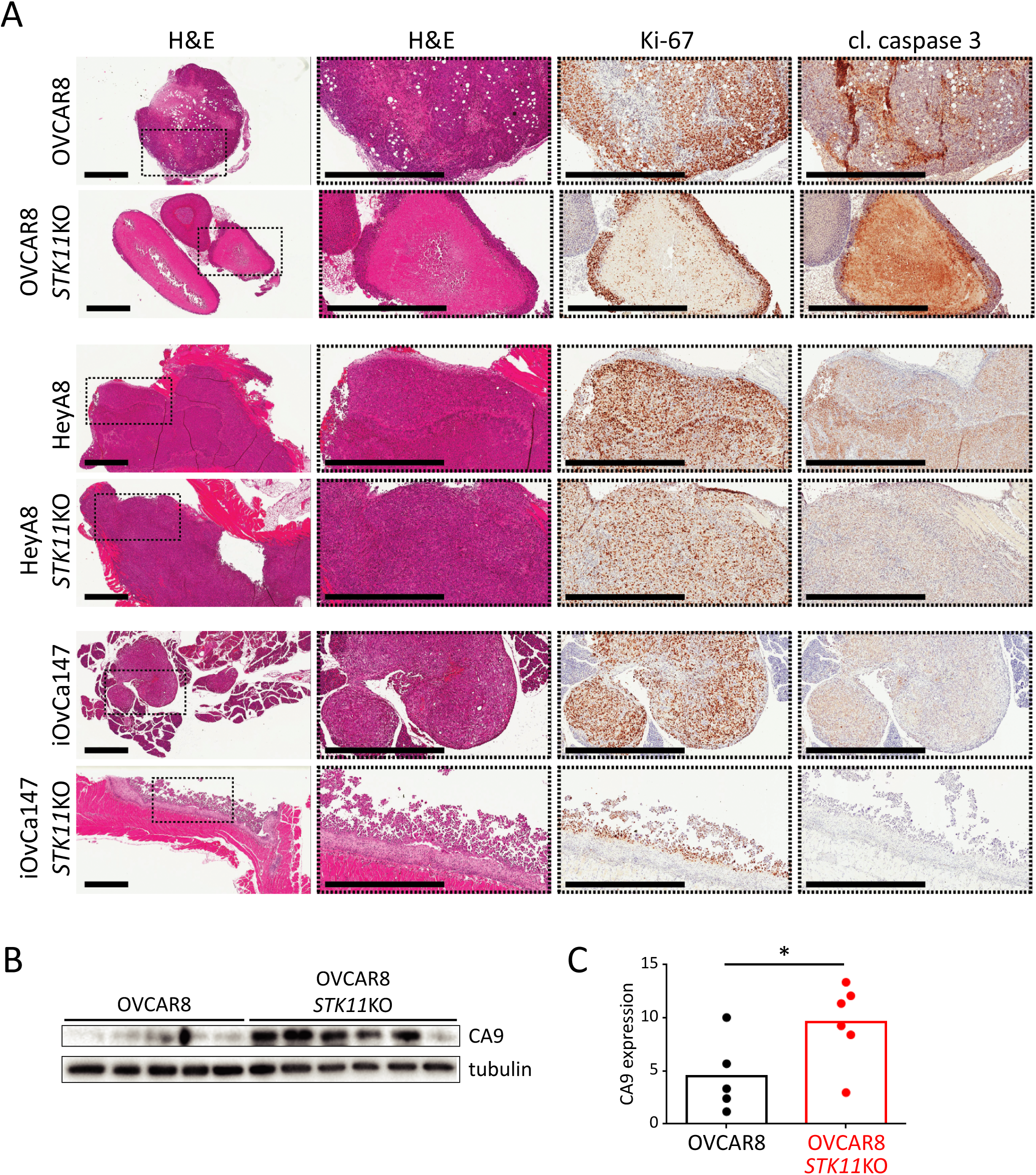
Histopathologic differences due to LKB1 loss in xenografted EOC tumours. **(A)** Serial sections were stained with hematoxylin and eosin (H&E), and immunostained using Ki67 for proliferation and cleaved caspase-3 for apoptosis. Boxes (dashed black lines) in the left-hand H&E panels at low magnification represent areas shown in the higher magnification panels to their right. Scale bars represent 1 mm. **(B)** Immunoblot analysis to determine relative carbonic anhydrase 9 (CA9) protein abundance as a marker of hypoxia in OVCAR8 and OVCAR8-*STK11*KO xenograft tumours. Each lane represents a tumour from a different mouse. **(C)** Densitometric analysis of immunoblot in (B). Data are presented as the pixel intensity volume for CA9 normalized to tubulin. Groups were compared using the two-tailed Student’s *t*-test (*, *p* < 0.05).

Notably, the significant decrease in *in vivo* metastatic potential for all three *STK11*KO EOC cell lines corresponded very well with our *in vitro* results using the spheroid model of metastasis (**Figure 2**). Furthermore, in cases where ascites was recovered from host mice, we observed the presence of multicellular aggregates that resembled the morphology of spheroids generated *in vitro* **(Supplemental Figure S3**). This indicates that spheroid biology in our *in vitro* model recapitulates the phenotype of spheroids formed in an established *in vivo* model of EOC metastasis.

### LKB1-independent AMPK phosphorylation in EOC cells, spheroids and xenograft tumours

We demonstrated previously that both LKB1 and phosphorylated AMPKα are coordinately upregulated in EOC spheroids [8], suggesting that LKB1-AMPK signaling is activated during EOC metastasis. Since LKB1 is commonly described as the major upstream kinase responsible for phosphorylation of AMPKα at Thr172 [9], we expected that phospho-AMPKα would be substantially decreased or ablated in *STK11*KO spheroids relative to parental controls. Instead, we found that phospho-AMPKα Thr172 was maintained in spheroids for all *STK11*KO cell lines tested (**Figure 5A**), indicating that in FTE and EOC spheroids AMPKα is phosphorylated by one or more other kinases independently of LKB1.

**Figure 5.**
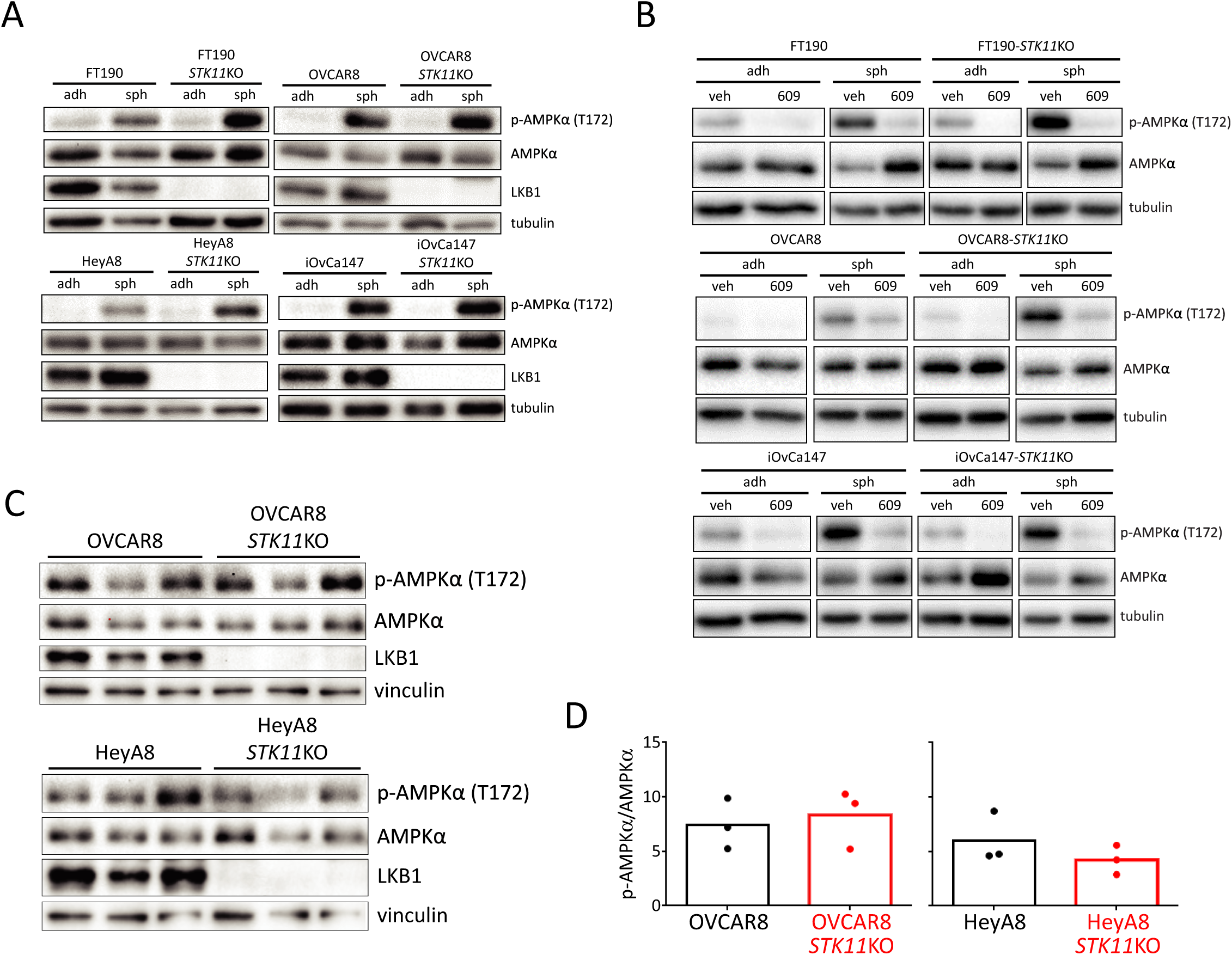
AMPKα phosphorylation is maintained in FTE and EOC *STK11*KO cells. **(A)** Immunoblot analysis to assess p-AMPKα (T172) in parental and *STK11*KO FTE and EOC cell lines cultured as adherent monolayers (adh) or spheroids in suspension (sph). LKB1 loss was confirmed, and tubulin was used as a loading control. **(B)** Immunoblot analysis to assess p-AMPKα (T172) in FTE, OVCAR8 and iOvCa147 cells cultured under adherent (adh) or suspension (sph) conditions in the presence of 10 μM STO-609 or the equivalent concentration (v/v) of DMSO (veh). **(C)** Immunoblot analysis to determine relative p-AMPKα (T172) levels in xenograft tumours. LKB1 loss was confirmed, and vinculin was used as a loading control. Each lane represents a tumour from a different mouse. **(D)** Densitometric analysis of immunoblots in (A). Data are presented as the pixel intensity volume for p-AMPKα (T172) normalized to AMPKα. Groups were compared using the two-tailed Student’s *t*-test with no significant difference observed.

Ca(2+)/calmodulin-dependent kinase kinase-beta (CAMKK2) is one principal alternative AMPK upstream kinase [33], and cell detachment can rapidly induce phospho-AMPKα at T172 in a CAMKK2-dependent manner in breast cancer cells [34]. Therefore, we treated FT190-*STK11*KO, OVCAR8-*STK11*KO, and iOvCa147-*STK11*KO spheroids and parental cell line controls with STO-609, a potent cell-permeable inhibitor of CAMKK2 [35]. STO-609 substantially decreased phospho-AMPKα in adherent cells and spheroids for all cell lines tested (**Figure 5B**), indicating that CAMKK2 is likely the major activating kinase for AMPKα in FTE and EOC cells, whether LKB1 is present or not. Importantly, analysis of tumour xenografts revealed that phospho-AMPKα was unaffected by loss of LKB1, also (**Figure 5C & D**). Indeed, LKB1 loss had no effect on OVCAR8 cells when subjected to several metabolic stressors— glucose deprivation, serum starvation, mitochondrial inhibition, and AMP mimetic treatment— which act through AMPK activity (**Supplemental Figure S4**). Taken together, these results imply that any of our observed phenotypic effects due to LKB1 loss in FTE and EOC cells and spheroids, and during EOC metastasis in mice, are most likely AMPK-independent.

### DUSP4 is upregulated due to sustained LKB1 loss

Since it appeared that AMPK was not mediating the phenotypic changes in EOC spheroids and tumours due to LKB1 loss, we sought to discover other potential effectors. To do so, we performed reverse phase protein array (RPPA) analysis comparing *STK11*KO knockout cells grown in adherent and spheroid culture using both OVCAR8 and HeyA8 cells. From this data, we observed distinct differences in protein expression between the OVCAR8 and HeyA8 spheroids due to LKB1 loss (**Figure 6A and Supplementary Table S2**), which may reflect the differences seen in spheroid culture and xenograft experiments between these two distinct EOC cell lines. However, one protein that was consistently increased due to LKB1 loss in both adherent cells and even more so in spheroids was dual-specificity phosphatase 4 (DUSP4). We observed a 1.86-2.06-fold increase in OVCAR8-*STK11* adherent cells and 1.76-2.57-fold increase in spheroid culture; in HeyA8-*STK11*KO cells, DUSP4 protein was elevated by 2.37-2.76-fold in adherent and 2.84-3.12-fold in spheroids. We confirmed this increase in DUSP4 protein expression among all three EOC cell lines where LKB1 was ablated (**Figure 6B**). In addition, DUSP4 was increased significantly in OVCAR8-*STK11*KO xenografts as compared with tumours derived from OVCAR8 cells (**Figure 6C and 6D**).

**Figure 6.**
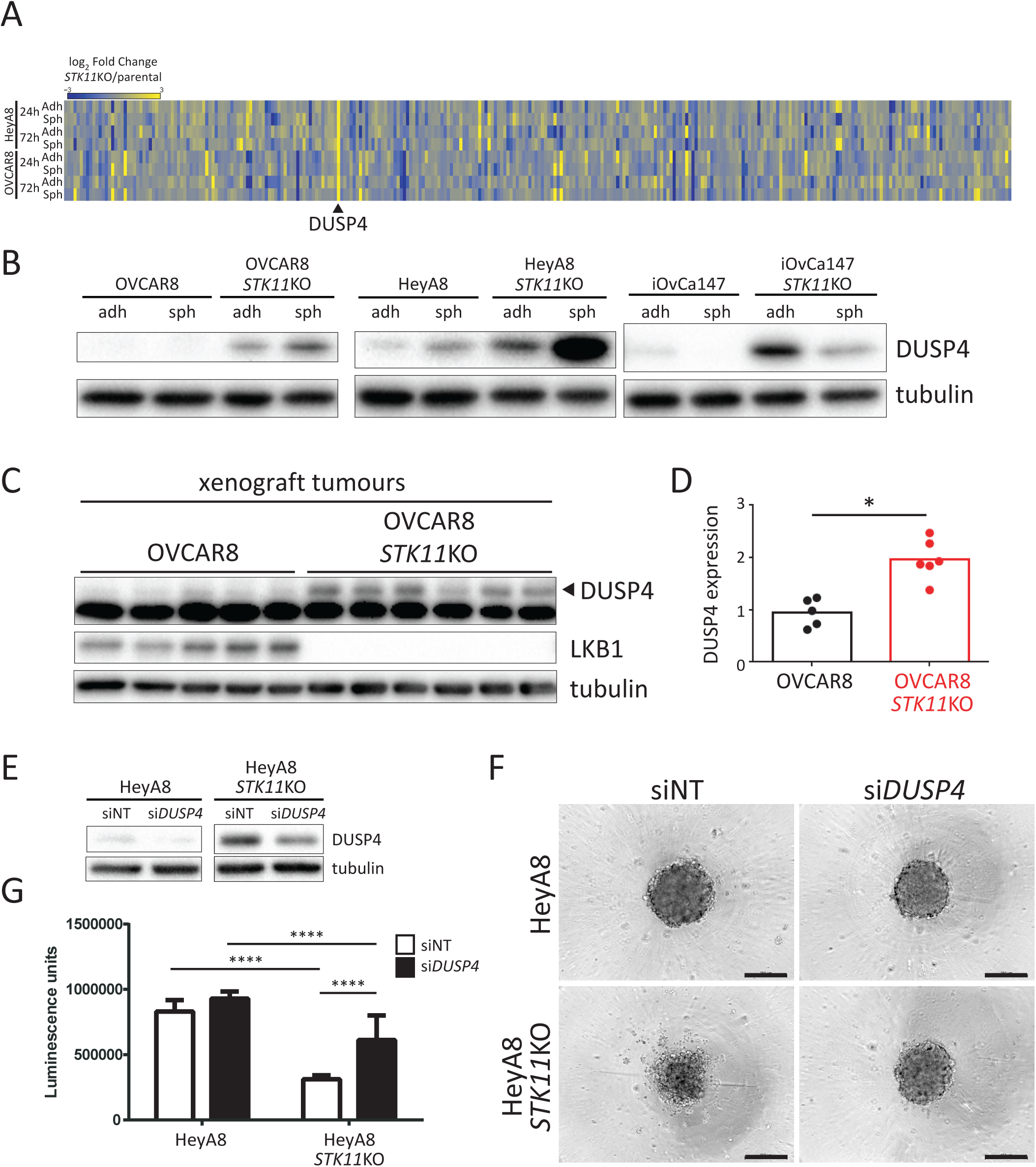
DUSP4 expression is increased due to sustained LKB1 loss. **(A)** Lysates from OVCAR8, OVCAR8-*STK11*KO, HeyA8 and HeyA8-*STK11*KO cells grown in adherent and spheroid culture were subjected to reverse-phase protein array analysis. Data is presented as a heat map of the log_2_-transformed fold-change in expression comparing *STK11*KO cells to their respective parental cell controls. DUSP4 expression was consistently increased due to LKB1 loss in every sample analyzed. **(B)** Immunoblot analysis confirming DUSP4 expression is increased in all three EOC cell lines lacking LKB1 as compared with their respective controls. **(C)** Immunoblot analysis demonstrating increased DUSP4 protein in OVCAR8-*STK11*KO tumour xenografts as compared with OVCAR8 tumours. LKB1 loss was confirmed, and tubulin used as a loading control. **(D)** Densitometric analysis of immunoblots in (C). Data are presented as the pixel intensity volume for DUSP4 normalized to tubulin with the mean value for OVCAR8 tumours set to 1. Statistical analysis was performed using a two-tailed Student’s *t-*test (*, *p* < 0.05). **(E)** Knockdown of *DUSP4* in HeyA8 and HeyA8-*STK11*KO cells as confirmed by immunoblot analysis. **(F)** Restoration of intact spheroid formation in HeyA8-*STK11*KO cells resulting from *DUSP4* knockdown. Scale bar represents 250 µm **(G)** Rescue of cell viability in HeyA8-*STK11*KO spheroids due to *DUSP4* knockdown as determined by Cell-Titer Glo assay in a 96-well ULA format. Two-way ANOVA and Tukey’s multiple comparisons test was performed (****, *p* < 0.0001).

DUSP4 is a dual-specificity phosphatase that typically targets several MAPKs to decrease their phosphorylation state and thus downstream signaling activity [36]. To this end, we performed immunoblot analysis of phosphorylated ERK, JNK and p38 among the three EOC cell lines lacking LKB1 as compared with controls. Interestingly, phospho-ERK1/2 was consistently decreased in EOC cells upon spheroid culture, however levels of phospho-p38 and phospho-JNK (p54 and p46) were either unchanged or different among the three cell lines (**Supplementary Figure S5**). Levels of phosphorylated ERK, JNK and p38, however, yielded no change due to LKB1 loss even though DUSP4 protein is universally increased in these cell lines. To further investigate whether upregulation of DUSP4 protein may impact the spheroid phenotype (i.e., reduced formation potential and cell viability) due to LKB1 loss, we performed transient *DUSP4* knockdown in HeyA8 and HeyA8-*STK11*KO cells (**Figure 6E**) and subsequently formed spheroids. We observed a restoration of intact spheroid formation by *DUSP4* knockdown in HeyA8-*STK11*KO spheroids (**Figure 6F**), as well as increased cell viability to levels similar to control HeyA8 spheroids (**Figure 6G**). Therefore, we speculate that DUSP4 is at least one protein mediating the reduced metastatic potential due to LKB1 loss in our *in vitro* spheroid model, and potentially decreased tumour formation and metastasis *in vivo*.

## DISCUSSION

Peritoneal metastasis in EOC is often associated with the accumulation of malignant ascites, which contains spheroids that play a key role in disease progression [3]. We previously reported that EOC cells require LKB1 to maintain spheroid viability and chemo-resistance [8]. Given the functional relevance of spheroids in EOC metastasis [3; 5-7], targeting LKB1 or its signaling partners in metastatic EOC may have therapeutic value. Herein, we now present additional evidence that EOC cells require LKB1 for maximal metastatic potential and add to the body of literature indicating that LKB1 can function in a pro-oncogenic capacity to promote disease progression. Based on our findings, and those of others, we propose that LKB1 or its AMPK-independent signaling partners may represent therapeutic vulnerabilities in metastatic EOC.

In the context of cancer, LKB1 has been commonly described as possessing tumour suppressor-like function. *STK11* is frequently deleted in non-small cell lung cancer (NSCLC) [37], and somatic nonsense mutations in *STK11* occur with high frequency in primary lung carcinomas and lung cancer cell lines [12], as well as in 20% of cervical cancer [13] and melanoma [11]. Loss of LKB1 in the ovarian surface epithelium has been linked to papillary serous ovarian cancer in a mouse model [18]. LKB1 protein expression is decreased in HGSOC relative to normal FTE tissue; however, this study also found that genomic loss of both *STK11* alleles was associated with improved patient survival [15]. In fact, our previous report [8] demonstrated that LKB1 expression is maintained in late-stage EOC and is required for cell survival in spheroids and promotes platinum resistance. Given that FTE cells are the most likely cell-of-origin in HGSOC, we had the direct ability to determine whether LKB1 may act as a tumour suppressor in FT190 cells with its loss resulting in their oncogenic transformation. However, we found that neither cell proliferation, nor colony formation, nor anchorage-independence, nor spheroid viability and growth were increased due to *STK11* gene inactivation. Since FT190 cells are immortalized by expression of SV40 large T antigen [30], our results also indicate that the combination of p53 and pRb inactivation with LKB1 loss is insufficient for malignant transformation of FTE cells. Given the report suggesting decreased LKB1 protein expression in pre-malignant STIC lesions and primary serous ovarian tumours [15], we propose that LKB1 re-expression and its pro-metastatic effects are manifest at later steps in ovarian tumorigenesis after initial neoplastic transformation.

A growing body of evidence suggests that LKB1 can function in context-dependent, pro-tumorigenic capacities. In line with its role in metabolism, deficiencies in LKB1-AMPK signaling sensitize some cancer cells to energy stress. LKB1-deficient NSCLC cells are more sensitive to the mitochondrial complex I inhibitor phenformin compared to NSCLC cells with intact LKB1 [38], and AMPK is required in A549 lung cancer cells for survival under glucose-deprived culture conditions [20]. However, LKB1 may also promote oncogenic potential in ways that are not directly linked to the energy stress response. Expression of LKB1 protein is associated with poor patient prognosis in hepatocellular carcinoma, and ectopic expression of LKB1 in Hep3B cells increases tumour xenograft growth [39]. Similar to our study of EOC cells, *STK11* knockdown decreases spheroid formation and clonogenicity of MCF-7 cells, indicating that LKB1 can enhance anchorage-independence and replicative potential in breast cancer cells [22]. We have not yet elucidated the mechanisms by which LKB1 promotes metastatic potential of EOC cells; however, our data provide additional evidence to support its pro-tumorigenic role in yet another cancer-specific context.

During metastasis, cancer cells must overcome significant hurdles to maintain viability and seed the formation of secondary tumours. Enhanced anoikis resistance is associated with greater aggressiveness and tumorigenicity in *in vivo* models of EOC [40]. We found that LKB1 loss had little effect on EOC cell growth under standard adherent culture conditions, while having a clear negative impact on EOC cell viability in suspension. Our findings imply that LKB1 supports anoikis resistance and anchorage-independence in EOC cells through mechanisms that do not involve AMPK. We previously reported that phosphorylation of AMPKα at Thr172, which is required for its full catalytic activity [41], is induced upon EOC spheroid formation [8]. Herein, we show that despite LKB1 loss, phospho-AMPKα levels in FTE and EOC spheroids remain elevated, which implies AMPKα activation occurs independently of LKB1 in our spheroid system. One such alternative kinase is CAMKK2 [33], which has been implicated in cell detachment-induced phosphorylation of AMPKα in several breast cancer cell lines [34]. Using STO-609, we provide evidence that AMPKα phosphorylation in FTE and EOC spheroids is largely mediated by CAMKK2. In fact, CAMKK2 inhibition using STO-609 does not affect EOC spheroid cell viability (Laski and Shepherd, unpublished), which corroborates our previous finding that knockdown of *PRKAA1/2* encoding AMPK catalytic subunits[8]. Thus, we conclude that phospho-AMPK is induced via CAMKK2 in EOC spheroids, but that AMPK activity is dispensable for maintaining spheroid cell viability in the context of LKB1 activity.

While AMPKα is often described as the primary cell stress signaling mediator downstream of LKB1, we demonstrate that essential LKB1 functions in EOC metastasis are AMPK-independent. LKB1 is known to phosphorylate at least 12 substrates that are similar in structure to AMPK, collectively termed AMPK-related kinases. Broadly, these kinases can regulate cell polarity, migration, and metabolism in specific cell contexts [42]. Salt-inducible kinases SIK2 and SIK3 may be required in HGSOC metastasis, but their dependence on LKB1 in this disease is unknown [43–45]. AMPK-related kinase ARK5/NUAK1 has been reported to be overexpressed in EOC and implicated in the regulation of EMT [46]. In line with this, NUAK1 expression in EOC is associated with poor patient prognosis [47]. Our group is currently pursuing additional NUAK1 functions as a substrate of LKB1 in metastatic EOC. Elucidation of these downstream mediators will likely be of clinical relevance since they may represent more specific and suitable therapeutic targets in metastatic EOC than LKB1 itself.

We postulate that EOC cells are adaptively reprogrammed to handle abrogated stress signaling due to sustained LKB1 loss. DUSP4 is commonly increased in NSCLC tumours possessing *STK11* loss-of-function mutations. Kaufman et al [48] identified an upregulated 16-gene expression signature, which included *DUSP4*, in LKB1-deficient NSCLC cells likely due to an altered oxidative stress response mediated by NRF2. We postulate that there is a similar dysregulated response to oxidative stress and hypoxia in the absence of LKB1 function in EOC spheroids and tumour xenografts. This could explain our observation of elevated hypoxia in OVCAR8-*STK11*KO xenograft tumours correlating with increased DUSP4 expression, perhaps resulting from chronic stress due to LKB1 loss in this cell line. It would be intriguing to study whether LKB1 signaling is required in response to tumour hypoxia and oxidative stress to regulate a downstream transcriptional network and mitigate these key metabolic stresses in cancer.

While LKB1 may act as a tumour suppressor in other cancers, we have provided additional evidence that LKB1 clearly possesses important functions in a pro-metastatic capacity during EOC progression. Consistent with the established notion that three-dimensional spheroids contribute to EOC dissemination, we have demonstrated that intact LKB1 is required in both *in vitro* spheroid and *in vivo* xenograft models of metastatic disease. Overall, we provide supportive evidence that LKB1 or its direct signaling partners represent viable therapeutic targets in metastatic EOC.

## MATERIALS AND METHODS

### Cultured cell lines

OVCAR8 and HeyA8 cell lines were cultured in RPMI-1640 (Wisent). FT190 and iOvCa147 cell lines were cultured in DMEM/F12 (Life Technologies). For all cell lines, growth medium was supplemented with 10% FBS. OVCAR8 and HeyA8 cells were obtained from the American Type Culture Collection and iOvCa147 cells were generated by us as described previously [23]. The immortalized human fallopian tube secretory epithelial cell line FT190 [30] was provided by R. Drapkin (University of Pennsylvania). All cell lines were authenticated by short tandem repeat analysis performed by The Centre for Applied Genomics (The Hospital for Sick Children, Toronto, ON, Canada), and routinely tested for mycoplasma using the Universal Mycoplasma Detection Kit (30-1012K, ATCC) or as performed by IDEXX BioResearch (Columbia, MO, USA) prior to xenograft experiments.

### Generation of *STK11*KO cell lines

The 20-nucleotide guide sequence targeting the *STK11* gene 5’-AGCTT GGCCC GCTTG CGGCG-3’ was selected using CRISPR Design Tool (http://tools.genome-engineering.org). Complementary oligonucleotides 5’-CACCG AGCTT GGCCC GCTTG CGGCG-3’ and 5’-AAACC GCCGC AAGCG GGCCA AGCTC-3’ (Sigma-Genosys) were annealed and ligated into the BbsI-digested restriction endonuclease site of pSpCas9(BB)-2A-Puro plasmid (gift from Dr. F. Dick, Western University) as per the protocol described in Ran et al. [49] to generate the pSpCas9-sgSTK11 plasmid. Cells were transfected with pSpCas9-sgSTK11 plasmid using LipofectAMINE 2000 (Invitrogen) according to manufacturer’s instructions, followed by treatment with 1 μg/mL puromycin for one day. After growth recovery, limiting dilution subcloning of potential *STK11*-knockout cells was performed and confirmation of *STK11* knockout by western blotting for LKB1 expression. A minimum of five clones lacking LKB1 protein expression were positively identified and subsequently mixed in equal ratios to generate the *STK11*KO cell line populations.

### Antibodies and reagents

Antibodies against LKB1 (#3050S), phospho-AMPKα (T172) (#2535S), AMPKα (#5832S), phospho-p44/42 ERK (T202/Y204) (#9101S), phospho-JNK (T183/Y185) (#4668P), phospho-p38 (T180/Y182) (#4511S), and DUSP4 (#5149S) were purchased from Cell Signaling (Danvers, MA) and used at 1:1000 in 5% bovine serum albumin (BSA)/TBS-T. Antibody against CA9 (AF2188; 1 μg/mL) was purchased from R&D Systems (Minneapolis, MN). Antibody against tubulin (T5168; 1:8000) and vinculin (V9264; 1:50 000) were purchased from Sigma (St. Louis, MO). Antibody against vinculin was purchased from HRP-conjugated antibodies against mouse IgG (NA931V; 1:10 000) and rabbit IgG (NA934V; 1:10 000) were purchased from Sigma. Immunohistochemistry was performed by Molecular Pathology (Robarts Research Institute, London ON, Canada); antibodies against Ki67 and cleaved caspase-3 were purchased from Abcam and used according to Molecular Pathology standard protocols. Oligomycin A (#11342) and AICAR (#10010241) were purchased from Cayman Chemical Company (Ann Arbor, MI, USA); STO-609 (#73862) was purchased from STEMCELL Technologies (Vancouver, BC, Canada).

### Immunoblot analysis

Isolation of protein lysates and western blotting were performed as described previously[8].

### Doubling time determination in adherent culture

Cells were seeded into 96-well adherent culture plates at a density of 1000 cells per well. Phase-contrast images were captured at 3-hour intervals for a total of 96 hours and confluence was measured using the IncuCyte Zoom imaging platform. Doubling time was determined by fitting an exponential growth curve to the confluence-over-time data (GraphPad Prism 6.05).

### Clonogenic assays

Cells were seeded at specific cell densities then fixed and stained when cells formed well-defined colonies with minimal overlap. OVCAR8 and OVCAR8-*STK11*KO cells: 250 cells/35-mm well, 8 days; iOvCa147 and iOvCa147-*STK11*KO cells: 1000 cells/35-mm well, 8 days; HeyA8 and HeyA8-*STK11*KO cells: 2500 cells/10-cm plate, 14 days; FT190 and FT190-*STK11*KO cells: 1000 cells/10-cm plate, 12 days. Cells were fixed and stained using the Protocol Hema 3 staining kit (Fisher Scientific). Colony number and size were determined using the *Fiji Is Just ImageJ* (Fiji) image processing software package [50]; the Trainable Weka Segmentation plugin [51] was used to generate classifiers to segment images into stained colonies and background. Images were converted into binary colour, “fill holes” and “watershed” functions were applied, and pixel-to-distance scale was set using the culture vessel diameter. Colony number and area were determined using the “analyze particles” function to outline each colony as a region of interest and provide colony counts.

### Soft agar assays

EOC cells were suspended at a density of 25 000 cells/1.5 mL in medium supplemented with 10% FBS and 1% melted soft agar; this suspension was added to each well containing solidified 2% agar. Additional growth medium was then added to each well, and plates were incubated for 14 days. Images of random fields of view were captured at 25x magnification using a Leica DMI 4000B inverted microscope. Colony number and size were determined using the Fiji image processing software package as described above.

### Spheroid viability assay

Cells were seeded into 24-well ULA cluster plates (50 000 cells per well in a volume of 1 mL). At the specified time points, the entire contents of each well were collected and trypsinized at 37°C with gentle agitation every 10 min until aggregates were no longer visible (10-30 min). Trypsin was inactivated and Trypan Blue Exclusion cell counting was performed using a TC10 Automated Cell Counter (Bio-Rad). For long-term spheroid growth assays, cells were seeded into a 96-well round-bottom ULA plate at a density of 2000 cells/well. Phase-contrast images were captured at 2-hour intervals for up to 12 days using the IncuCyte Zoom imaging platform. Single spheroids were isolated at specific time points post-seeding (3, 7, 14, and 21 days) and transferred into separate wells of a 48-well adherent culture plate, incubated for 2 days, then attached spheroids were fixed and stained using the Protocol Hema 3 staining kit (Fisher Scientific).

### Xenotransplantation assays

NOD/SCID female mice (8-10 weeks old; Charles River) were inoculated by intraperitoneal injection with 150 µL of PBS containing the following numbers of cells: OVCAR8/OVCAR8-*STK11*KO, 4 × 10^6^; HeyA8/HeyA8-*STK11*KO, 1 × 10^6^; iOvCa147/iOvCa147-*STK11*KO, 2 × 10^6^. For survival analyses (OVCAR8, OVCAR8-*STK11*KO, HeyA8, HeyA8-*STK11*KO, iOvCa147 and iOvCa147-*STK11*KO cells), mice were monitored daily after intraperitoneal injection and euthanized using standard criteria for humane endpoints (i.e., lethargy, hunched posture, impaired breathing, extreme weight loss, excessive ascites). Mice were provided chow (cat. no. 2919, Envigo) and water *ad libitum* throughout the study. All animal experiments were approved by Institutional Animal Care and Use Committee of the University of Western Ontario and carried out in accordance with the approved guidelines.

### Inhibition of CAMKK2

Cells were seeded into a 6-well adherent culture plate, and treatment was initiated the following day with 10 μM STO-609 or an equivalent volume of DMSO. For analysis of spheroids, cells were seeded into a 6-well ULA plate and treated with 10 μM STO-609 or DMSO at the time of seeding. Lysates were generated for immunoblot analysis at 24 h (FT190, FT190-*STK11*KO, iOvCa147 and iOvCa147-*STK11*KO cells) or 72 h (OVCAR8 and OVCAR8-*STK11*KO cells).

### Induction of metabolic stress

For glucose deprivation experiments, cells were seeded into 96-well adherent culture plates (2500 cells/well) in complete growth medium. The following day, growth medium was aspirated, washed once with serum- and glucose-free medium, and replaced with medium containing 10% dialyzed FBS (Invitrogen) and 0.1 g/L glucose. For serum deprivation experiments, cells were incubated with medium containing 0.25% FBS. For inhibition of mitochondrial complex V using oligomycin A or pharmacological activation of AMPKα using AICAR, cells were seeded into 96-well adherent culture plates (1000 cells/well) in complete growth medium in a volume of 150 μL. The following day, 50 μL of complete growth medium containing oligomycin, AICAR, or vehicle (DMSO) was added to each well. Viability was assessed 72 h later using the CyQuant Cell Proliferation assay as per manufacturer’s instructions.

### Reverse phase protein array (RPPA) analysis

Adherent cells were harvested at 24 h and 72 h post-seeding by trypsinization followed by addition of RPMI with 10% FBS; spheroids were directly collected from ULA plates at these same time points. Cells were pelleted (800xg, 4°C) and washed twice in ice cold PBS, then flash-frozen on dry ice, and stored at −80°C until sample submission. RPPA analysis was performed by the Functional Proteomics RPPA Core Facility at the MD Anderson Cancer Center according to their standard protocol [https://www.mdanderson.org/research/research-resources/core-facilities/functional-proteomics-rppa-core/antibody-information-and-protocols.html].

Normalized data and log2-fold changes are summarized in **Supplementary Table S2**. A heat map was generated using log2-fold change data and the Matrix2png interface (https://matrix2png.msl.ubc.ca).

### *DUSP4* knockdown

Transfections were performed of cells seeded in 6-well plates using DharmaFECT1 as per manufacturer’s protocol (Dharmacon; Thermo Fisher Scientific Inc., Waltham, MA). *DUSP4* (M-003963-03) and non-targeting control pool #2 (D-001206-14-05) siGENOME SMARTpool siRNA was used. After 72 h, trypsinized cells were counted and seeded at a density of 5 × 10^4^ cells per well in 24-well ULA cluster plates and 2 × 10^3^ cells per well in 96-well round-bottom ULA cluster plates. Cell viability was determined at 72 h post-seeding in 96-well round-bottom ULA cluster plates using CellTiter-Glo® (Promega). Western blot analysis was performed on adherent cell lysates at 72 h post-seeding to confirm *DUSP4* knockdown.

### Statistical analysis

All statistical analyses were performed using GraphPad Prism 6.05 (GraphPad Software, San Diego, CA). The results from *in vitro* analyses were assessed using a two-tailed Student’s *t*-test or two-way ANOVA with Tukey’s multiple comparisons test. Tumour latency analysis in mice was tested for significance using the log-rank test. In all cases, a *p*-value of less than 0.05 was considered statistically significant.

## Supporting information

Supplementary Video HeyA8 spheroid

Supplementary Video HeyA8-STK11KO spheroid

Supplementary Video OVCAR8 spheroid

Supplementary Video OVCAR8-STK11KO spheroid

## ACKNOWLEDGEMENTS

We are grateful to Ronny Drapkin for offering the use of the FT190 cell line and Frederick Dick for the pSpCas9(BB)-2A-puro plasmid. We thank Rene Figueredo for performing the tumour cell injections into mice, and Yingke Liang and Yixin Jiang for assisting with data analysis. This research was supported by funding from the Canadian Institutes for Health Research (Grant No. 136836) (TGS) and a London Regional Cancer Program Catalyst Grant (TGS) with funds from the London Run for Ovarian Cancer. We are also very grateful to the many donors to the Mary and John Knight Translational Ovarian Cancer Research Unit through the London Health Sciences Foundation for additional infrastructure funding, including the Leica DMI 4000B inverted microscope and IncuCyte ZOOM imaging system.

## AUTHOR CONTRIBUTIONS

GED and TGS conceived of the project. AB, YRV and TGS carried out experiments. AB, YRV, GED and TGS interpreted experimental data. AB and TGS wrote the manuscript, and GED assisted with editing.

