## Supplementary Figures for "AMPK-independent LKB1 activity is required for efficient epithelial ovarian cancer metastasis"

### Supplementary Figure S1

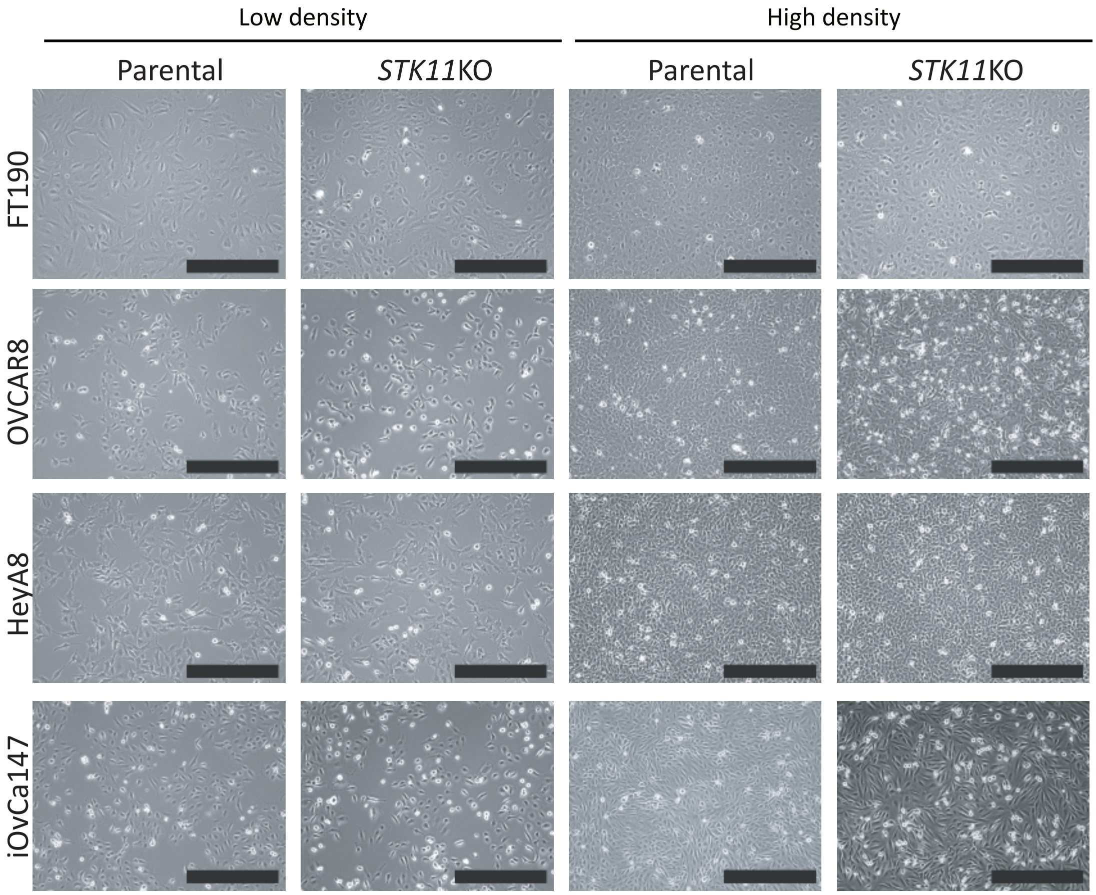


**Supplementary Figure S1. Morphology of *STK11*KO cell lines in adherent culture.** Images were captured at low and high cell densities. Scale bars indicate 500 µm.

### Supplementary Figure S2

**A**

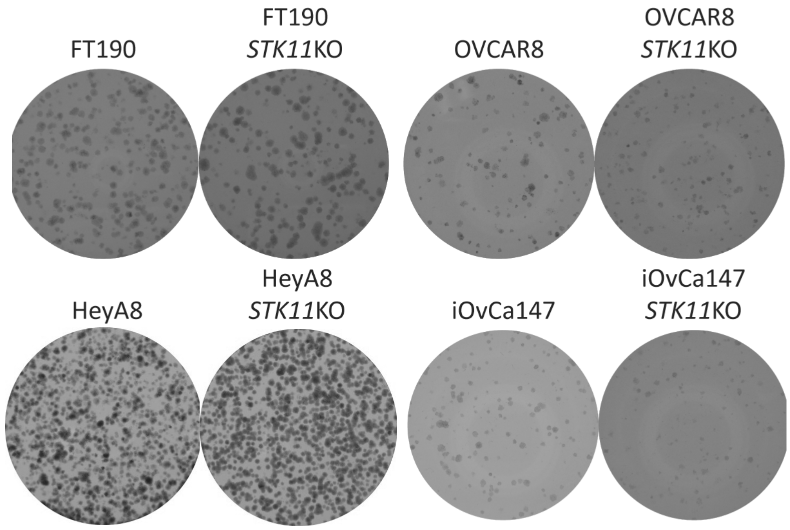


**B**

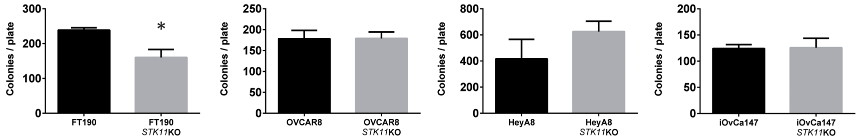


**Supplementary Figure S2. Effect of LKB1 loss on cell clonogenicity.** (A) Representative images of Hema-3 stained colony formation assays, performed as described in Materials and Methods. (B) Colonies were counted automatically using the Trainable Weka Segmentation plugin (version 3.2.27) in the Fiji software package. Data are presented as the number of colonies per dish or well (mean; n=3 for each group). Parental and *STK11*KO groups were compared using the two-tailed Student's *t*-test (*, $p < 0.05$).

#### Supplementary Fig S3

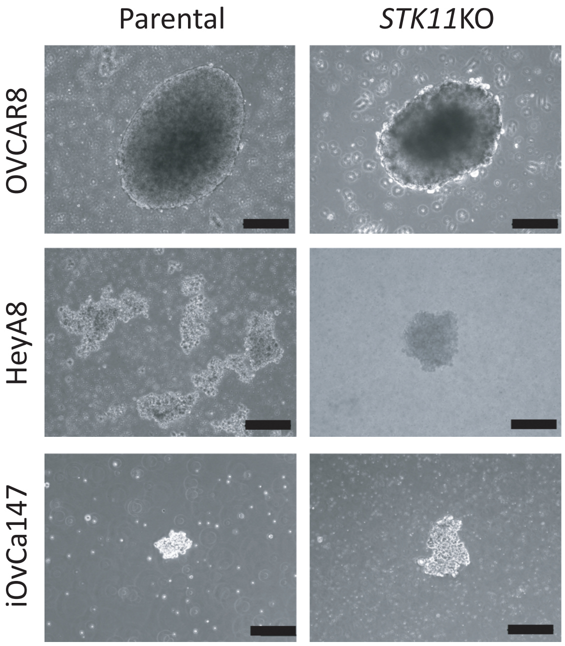


**Supplementary Figure S3. Spheroids isolated from tumour cell mouse xenograft ascites.** Ascites was removed at the time of necropsy, placed into adherent culture dishes containing complete growth medium, and phase-contrast images were captured immediately thereafter. Scale bars indicate 200 µm.

#### Supplementary Figure S4

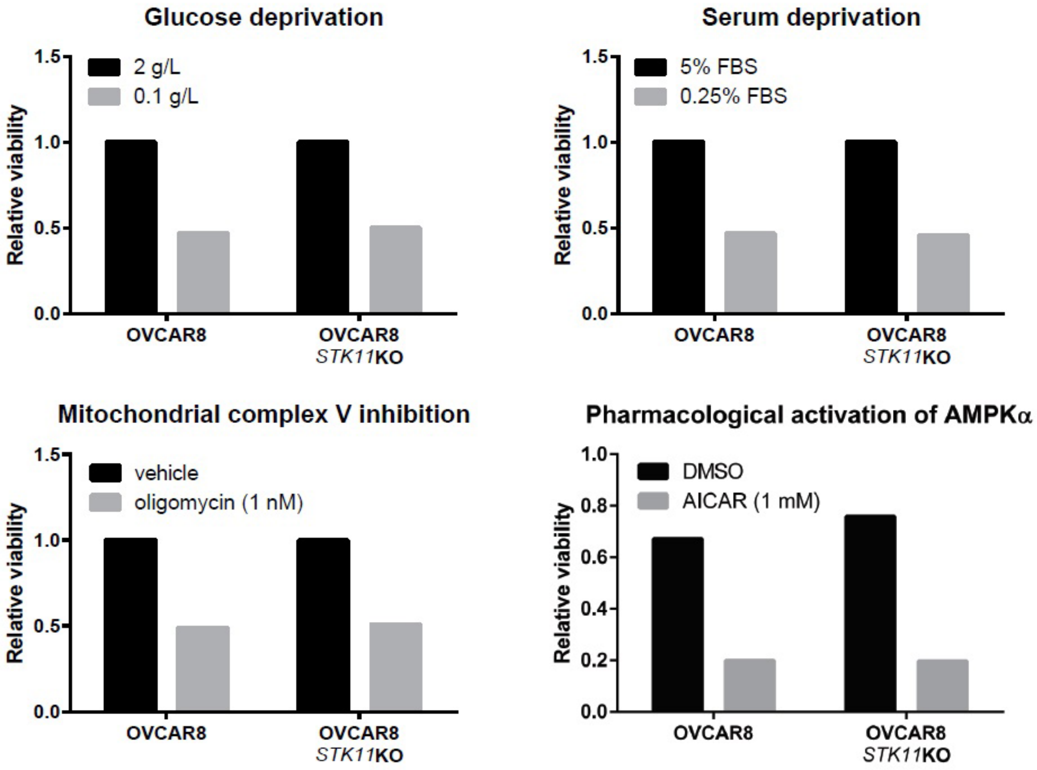


**Supplementary Figure S4. Loss of LKB1 does not alter sensitivity of OVCAR8 cells to several metabolic stresses.** Cells were seeded and treated the following day as described in the Materials and Methods. Relative cell viability was determined after 3 days using the CyQuant Cell Proliferation Assay (Promega).

### Supplementary Figure S5

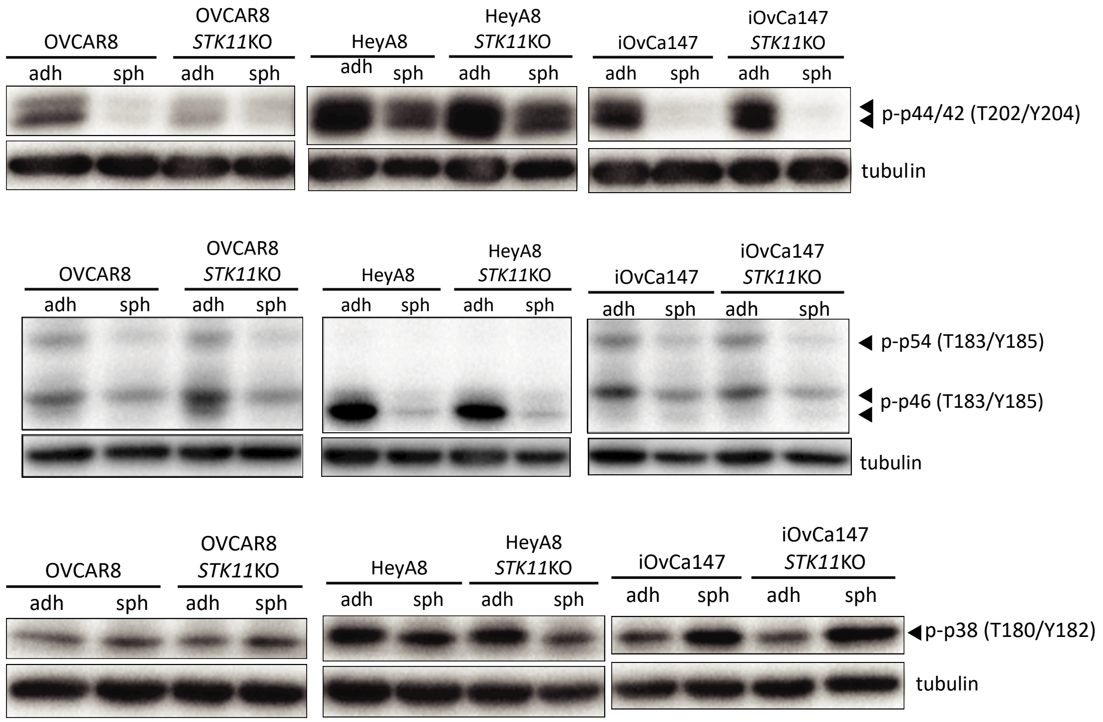


**Supplementary Figure S5. LKB1 loss does not alter MAPK phosphorylation in EOC cells and spheroids.** Immunoblot analysis of phosphorylated ERK (p44/42), JNK (p54/p46), and p38 in OVCAR8, OVCAR8-*STK11*KO, HeyA8, HeyA8-*STK11*KO, iOvCa147, and iOvCa147-*STK11*KO cells grown in adherent (adh) and spheroid (sph) culture.
